# Immuno-Detection by sequencing (ID-seq) enables large-scale high-dimensional phenotyping in cells

**DOI:** 10.1101/158139

**Authors:** Jessie A.G. van Buggenum, Jan P. Gerlach, Sabine E.J. Tanis, Mark Hogeweg, Jesse Middelwijk, Ruud van der Steen, Cornelis A. Albers, Klaas W. Mulder

## Abstract

Cell-based small molecule screening is an effective strategy leading to new medicines. Scientists in the pharmaceutical industry as well as in academia have made tremendous progress in developing both large-scale and smaller-scale screening assays. However, an accessible and universal technology for measuring large numbers of molecular and cellular phenotypes in many samples in parallel is not available. Here, we present the Immuno-Detection by sequencing (ID-seq) technology that combines antibody-based protein detection and DNA-sequencing via DNA-tagged antibodies. We used ID-seq to simultaneously measure 84 (phospho-)proteins in hundreds of samples and screen the effects of ~300 kinase inhibitor probes on primary human epidermal stem cells to characterise the role of 225 kinases. Our work highlighted a previously unrecognized downregulation of mTOR signaling during differentiation and uncovered 13 kinases regulating epidermal renewal through distinct mechanisms.

## Introduction

Quantification of protein levels and phosphorylation events is central to investigating the cellular response to perturbations such as drug treatment or genetic defects. This is particularly important for cell-based phenotypic screens to discover novel drug leads in the pharmaceutical industry. However, the complexity of biological and disease processes is not easily captured by changes in individual markers. Currently, a major limitation is the trade-off between the number of samples and the number of (phospho-)proteins that can be measured in a single experiment. For instance, immunohistochemistry^1^ and immunofluorescence^2^ allow high-throughput protein measurements using fluorescently labelled antibodies. However, these methods are limited in the number of (phospho-)proteins that can be measured simultaneously in each sample due to spectral overlap of the fluorescent reporter dyes. One commercial solution, Luminex®, has circumvented this limitation by using color barcoded antibody loaded beads and allows multiplexing of some 50 proteins per sample^3^–^5^. However, this approach requires cell lysis and does currently not include phospho-specific signaling detection. Several alternative approaches based on antibody-DNA conjugates have been developed in recent years^6^,^7^. For instance, Ullal et al. use the Nanostring system to quantify 88 antibody-ssDNA oligo conjugates in fine needle aspirates^6^. Although powerful, this strategy is not well suited for high-throughput applications. Furthermore, the commercial Proseek^®^ strategy entails a proximity extension assay using pairs of ssDNA oligo coupled antibodies in combination with quantitative PCR as a read-out^7^. This assay is generally performed on cell lysates and currently there are no assays for phospho-proteins available. In addition, several other recently described antibody-DNA conjugate based methods that use high-throughput sequencing as a read-out detect only a few epitopes or at low sample throughput^8–12^, limiting their scope. Here, we present Immuno-Detection by sequencing (ID-seq) as a streamlined universal technology for measuring large numbers of molecular and cellular phenotypes in many samples in parallel. We show that high-throughput sequencing of antibody coupled DNA-barcodes allows accurate and reproducible quantification of 84 (phospho-)proteins in hundreds of samples simultaneously. We applied ID-seq in conjunction with the Published Kinase Inhibitor Set (PKIS) to start investigating the role of >200 kinases in primary human epidermal stem cell renewal and differentiation. This demonstrated a previously unrecognized downregulation of mTOR signaling during differentiation and uncovered 13 kinases regulating epidermal renewal through distinct mechanisms.

## Results

### Precise and sensitive multiplexed (phospho-)protein detection via sequencing antibody-coupled DNA-tags

We designed the Immuno-Detection by sequencing (ID-seq) technology to simultaneously measure many proteins and post-translational modifications in high-throughput (Figure 1a). At the basis of ID-seq lie antibodies that are labelled with a double-stranded DNA-tag^13^ containing a 10 nucleotide antibody-dedicated barcode and a 15 nucleotide Unique Molecular Identifier (UMI, Figure S1, Supplementary note 1). Each antibody signal is now digitized and non-overlapping, allowing many antibodies to be combined and measured simultaneously. Following immunostaining and washing, DNA-barcodes are released from the antibodies through reduction of a chemically cleavable linker^13^ and a sample-specific barcode is added through PCR. Finally, samples are pooled to prepare an indexed sequencing library (Figures 1a, S1 and Supplementary note 1). This triple barcoding strategy facilitates straightforward incorporation of hundreds (and potentially thousands) of samples per experiment and achieves count-based quantification (Figure S2 and Supplementary note 2) over four orders of magnitude (Figure S3). Furthermore, analyses of 17 antibody-DNA conjugates using singleplex and multiplexed measurements show high correspondence (R = 0.98 ± 0.046), demonstrating that multiplexing does not interfere with antibody detection (Figure 1b). Moreover, the ID-seq library preparation procedure is reproducible (R=0.98, Figure 1c) and precise, as determined using nine distinct DNA-tag sequences per antibody, serving as technical replicates (R > 0.99, Figure S4). Thus, the ID-seq technology allows precise and sensitive multiplexed protein quantification through sequencing antibody-coupled DNA-tags.

**Figure 1.**
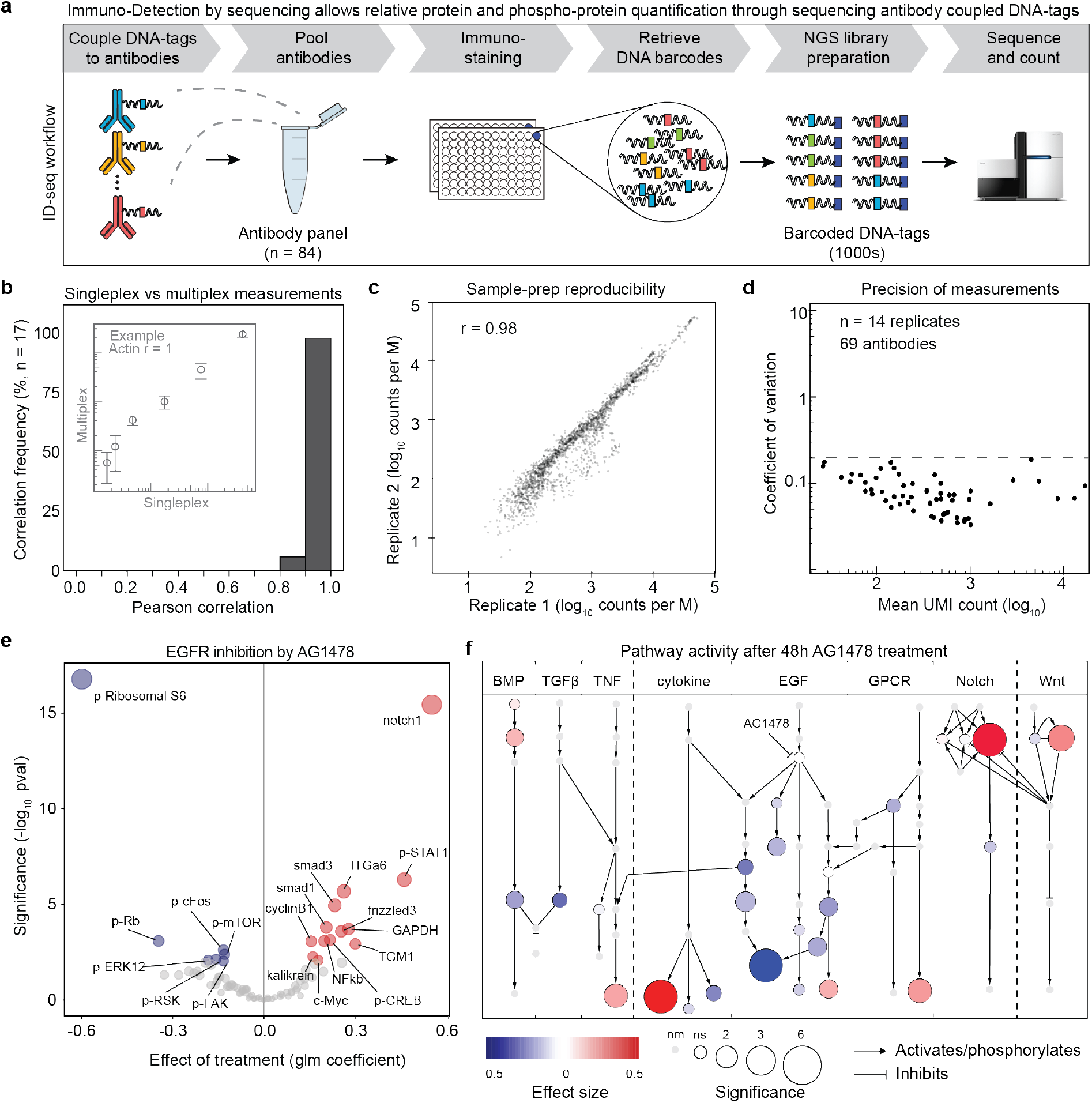
Immuno-Detection by Sequencing (ID-seq) technology development. (**a**) Concept of the ID-seq technology. First, pool DNA-tagged antibodies. Second, perform multiplexed immunostaining on fixed cell populations, and release DNA-tags. Third, barcode the released DNA-tags through a 2-step PCR protocol. Finally, sequence the barcoded DNA-tags via next generation sequencing (NGS) and count barcodes. (**b**) Signals from singleplex epitope detection (via an immuno-PCR measurement) were compared with multiplexed epitope detection using ID-seq. The histogram summarises correlations between 17 immuno-PCR and corresponding ID-seq measurements. Insert panel illustrates an example from the Actin antibody showing signal mean and sd (n=4). Underlying data for all 17 antibodies can be found in Figure S4. (**c**) Scatter plot indicates the high reproducibility of PCR-based ID-seq library preparation (r=Pearson correlation). Libraries from the same released material were prepared on separate occasions and analysed in different sequencing runs. (**d**) The scatterplot shows the counts (mean) and coefficient of variation from 69 antibody-DNA conjugates (n=14 biological replicates). Dashed line indicates 20% variation. (**e**) Volcano plot shows the effect (estimate) and significance (-log1_0_ pval) of AG1478 treatment (n=6), based on the model analysis of ID-seq counts (Supplementary Note 3). Significance (-log1_0_ pval) determines node size. Red nodes show significantly increased (p < 0.01) and blue nodes show significantly decreased (p < 0.01) (phosphor-)protein levels. (**f**) Pathway overview of ID-seq measurements after 48 hours of AG1478 treatment. Colour indicates the effect size, and node size represents the significance of effect (-log1_0_ pval). Light grey nodes without border indicate not measured (nm) proteins (see Supplementary Figure 7 for (phospho-)protein identities).

### Constructing a 70 antibody-DNA conjugate ID-seq panel

In order to fully exploit the multiplexing capacity of ID-seq we obtained 111 antibodies targeting intracellular and extracellular epitopes. All of the selected antibodies were validated for specificity in immunofluorescence (IF), immunohistochemistry (IHC) and/or fluorescence activated cell-sorting (FACS) applications by the vendor (see supplementary table 1 for details and links to datasheets). As these applications include cell fixation we reasoned that this selection would increase the chance of identifying antibodies that are suitable for ID-seq. From these 111 antibodies, 84 showed robust signals in In-Cell-Western /IF and/or immuno-PCR experiments using antibody dilutions and/or IgG control antibodies. Next, to increase our confidence in this set of antibodies, we performed a series of experiments using IF, immuno-PCR and/or ID-seq as a read-out to verify that the tested antibodies show the expected signal dynamics in response to specific perturbations. These perturbations included: induction of differentiation, stimulation with epidermal growth factor (EGF) or bone morphogenetic proteins (BMP), induction of DNA damage signalling with mitomycin C or hydroxyurea, as well as inhibition of EGF and BMP signalling with the small molecule inhibitors AG1478 and DMH1, respectively. In addition, signals of a subset of phospho-specific antibodies were decreased upon phosphatase treatment of fixed cell populations. Taken together, 64 out of the 84 antibodies exhibited the expected protein or phospho-protein dynamics in our primary skin stem cells in these experiments, whereas the rest was stable (Supplementary table 1 and Figure S5), indicating their utility in ID-seq. These 84 antibodies displayed ~75-fold signal over no-cell background, a measure of technical noise (Figure S6). Moreover, we found that the variability of the signals from a subpanel of 69 antibody-DNA conjugates was below 20% among 14 biological replicates (coefficient of variation, CV < 0.2, Figure 1d), demonstrating the precision and reproducibility of highly multiplexed ID-seq measurements. Taken together, these experiments enabled us to construct a panel of 70 antibody-DNA conjugates to evaluate protein levels and intracellular signalling of fixed primary human skin stem cells. This panel covers a broad range of biological processes including cell cycle, apoptosis, DNA damage, epidermal self-renewal and differentiation, as well as intracellular signalling status for the EGF, G-protein coupled receptors, calcium signalling, TNFα, TGFβ, Notch, BMP and Wnt pathways (Figure S7a and supplementary table 1). Of note, the nature of the selected and validated antibodies should make this panel broadly applicable to many other human (and mouse) derived cell systems.

### Exploring (phospho-)protein dynamics in human epidermal stem cells with ID-seq

Primary human epidermal stem cells (keratinocytes) depend on active epidermal growth factor receptor (EGFR) signalling for self-renewal *in vitro* and *vivo*^14^. We inhibited this pathway using the potent and selective inhibitor AG1478 (10 μM, 48 hrs) to determine whether ID-Seq recapitulates keratinocyte biology. In these experiments, we would expect to at least observe dynamic changes in downstream EGFR signalling pathway activity, as well as in the expression of differentiation-associated proteins. To analyse the effects of AG1478 treatment on each antibody signal, we developed a generalised linear mixed (glm) model that takes into account the negative binomial distribution of ID-seq count data and incorporates potential sources of variation (e.g., replicates, batches, sequencing depth). This model derives the effect (‘estimate’) of treatment on each antibody, followed by a likelihood ratio-test to determine the significance of the effect (Supplementary note 3). We identified 13 increased and 7 decreased (phospho-)proteins upon AG1478 treatment (p < 0.01, Figure 1e). Up-regulation of the known differentiation markers transglutaminase 1 (TGM1) and Notch1 confirmed successful differentiation. Next, we projected the estimates and significance levels of our ID-seq results onto a literature derived signalling network (Figure 1f, see Figure S7a for node identities). As expected, EGFR pathway activity was down regulated upon AG1478 treatment. We also identified effects on the activity of several other pathways, including the bone morphogenetic protein (BMP) and Notch cascades, which are known players in epidermal biology^15–18^. RT-qPCR analysis revealed that these effects arose from changes in mRNA expression of BMP ligands and Notch receptors (Figure S7b). We confirmed activation of the BMP and Notch pathways by RT-qPCR analysis of their classical downstream target genes Id2 and Hes2, respectively (Figure S7b). These results demonstrate the potential of ID-seq and our glm model to distinguish different treatment conditions by quantifying changes in (phospho-)protein dynamics.

### Identification of kinase inhibitor probes affecting epidermal stem cell differentiation

Extracellular signals involved in epidermal renewal and differentiation are widely studied and include EGF, TGFβ, BMP, Notch ligands and Wnts^15,19–23^. However, the contributions of different intracellular effector kinases on renewal and differentiation are not well documented. To start addressing this issue, we applied ID-seq to human epidermal keratinocytes treated with the Published Kinase Inhibitor Set (PKIS), an open-source chemical probe library^24–26^ containing ~300 small molecules targeting 225 kinases across all major kinase families in the human proteome (Figure 2a). To determine the effects of kinase inhibition at the molecular level, cells were seeded in 384-well plates, treated with 294 PKIS compounds for 24 hours and subjected to ID-seq with a panel of 70 antibodies. Replicate screens were highly correlated (R = 0.98) and had low UMI duplicate rates (1.2%), indicating high data quality (Figure S8). We annotated the effect and its significance for each kinase inhibitor probe on each of the measured molecular phenotypes (as measured by our antibody conjugates) and used the results of this analysis to interrogate the effect of kinase inhibition on skin stem cell biology.

**Figure 2.**
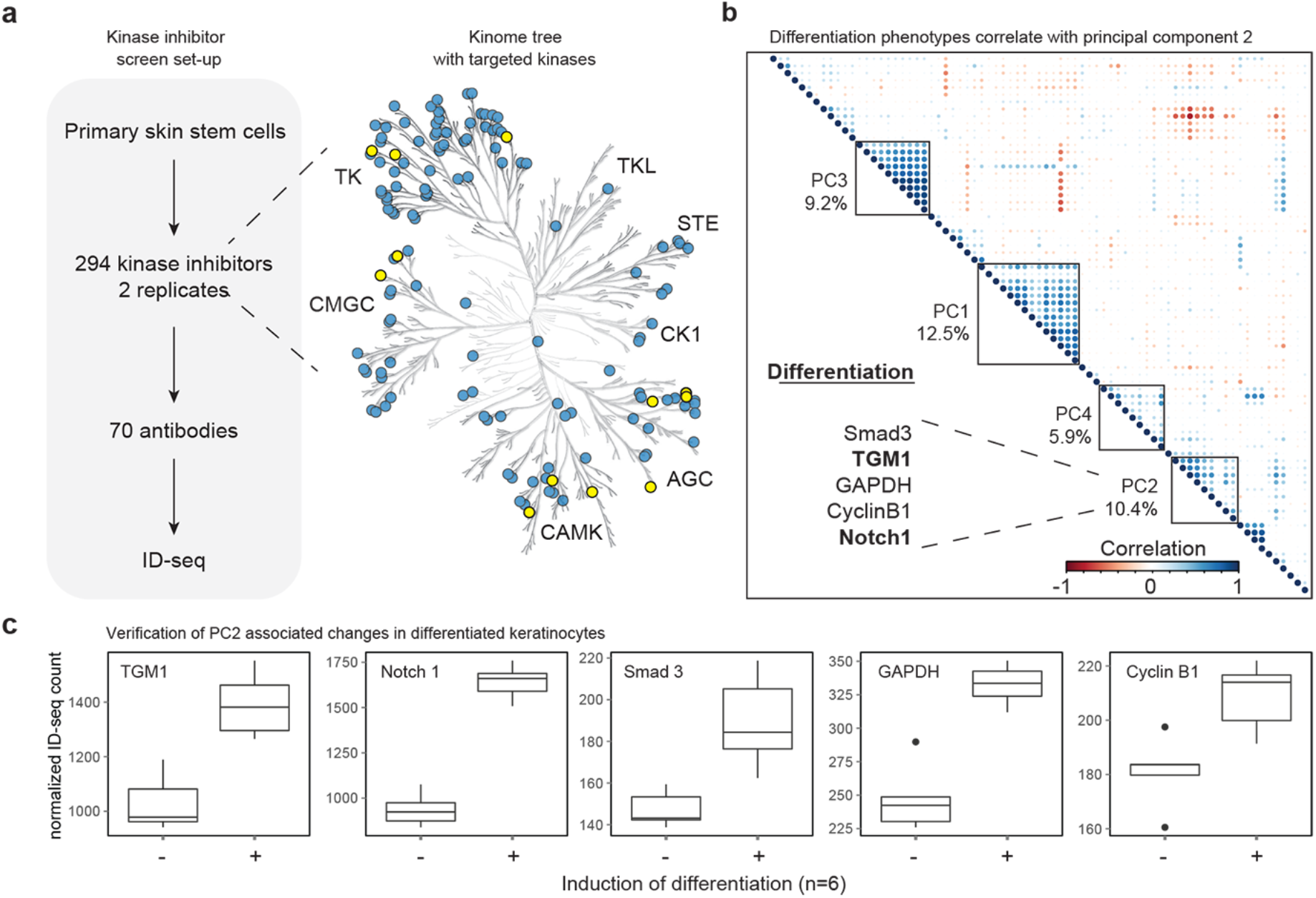
ID-seq screen of Published Kinase Inhibitor Set (PKIS) identifies probes inducing skin stem cell differentiation. (**a**) Schematic overview of screen set-up. Kinase-tree shows kinases targeted by the inhibitory probes (blue) and significantly enriched kinases (yellow, Figure 4a). The probes target all major kinase families (TKL, STE, CK, AGC, CAMK, CMGC and TK). (**b**) To combine PKIS probe effects on multiple ID-seq phenotypes to one measure, we performed principal component analysis on the PKIS dataset using the signed log_10_ p-values of the ID-seq analysis. Then, we clustered all phenotypes and the top 5 PCs to identify the PC summarising differentiation of the skin stem cells (in bold phenotypes TGM1 and Notch1). (**c**) ID-seq measurement of differentiation marker TGM1, Notch *1*, GAPDH, CyclinB1 and Smad3 upon inhibition of EGFR shows differentiation-induced phenotypic changes.

A key advantage of the high multiplexing capacity of ID-seq is its potential to measure multiple antibodies reflecting a given biological process simultaneously. We anticipate this to result in a more reliable and comprehensive measurement of the affected processes than quantification of a single marker. We exploited the multiplexed nature of the ID-seq data by combining the individual phenotypic ID-seq measurements into principal components (PCs) through principal component analysis. This essentially aggregates the phenotypes that jointly explain independent fractions of variation in the data into a single score that represents the effect of the inhibitory probes on the skin stem cells. To determine the underlying processes associated with each PC per probe, we correlated and clustered the measured phenotypes with the top 4 principal components explaining 38% of total variation in the dataset (Figures 2b, S9). As expected from a screen using kinase inhibitor probes a considerable fraction of this variation is associated with effects on signalling pathway activity phenotypes, as represented by PC1 (Figure 2b). The second largest principal component, PC2, strongly correlates with proteins that are significantly up-regulated upon differentiation, including the known marker proteins TGM1 and Notch1 (Figures 2b, ^15^,^27^). Interestingly, Cyclin B1, GAPDH and Smad3 were also included in this cluster. We confirmed that these phenotypes truly reflect keratinocyte differentiation, by forcing the cells to differentiate using the EGFR inhibitor AG1478 for 48 hours and subjecting these samples to ID-seq (n=6). Indeed, protein levels of TGM1, Notch1, Smad3, CyclinB1 and GAPDH are upregulated upon differentiation, corroborating the results of our screen (Figure 2c). To validate this further, we selected 18 probes that showed high PC2-scores from the PKIS library for colony formation experiments, the gold standard *in vitro* assay for epidermal stem cell activity (Figure S11a,b, ^28^). Automated image analysis was used to quantify colony number, colony size (and size distribution), as well as the level of the differentiation marker TGM1 level per colony for each probe (n = 3 replicates). Fifteen out of the eighteen tested probes showed a significant effect on at least one of the measured colony phenotypes (Figure S11a,b). This indicates that that high-PC2 probes indeed affect epidermal cell colony forming capacity and authenticates the PC2-score as a *bona fide* reflection of differentiation.

### Processes distinguishing differentiating and renewing epidermal stem cells

We gathered that the top and bottom 10% of PC2-ranked probes are likely to distinguish the differentiating (high-PC2) and non-differentiated (low-PC2) epidermal cell states. To determine which cellular processes are different between these two cell states, we determined the molecular phenotypes that display a significant in- or decrease (p < 0.01, 1% FDR) between cell-populations with high-PC2 versus low-PC2. This revealed involvement of the Wnt pathway (measured by Fzd3 and phosphorylated-LRP6), MAPK signalling (phospho-p38, phospho-SRC, phospho-cFOS and phospho-RSK), integrin-mediated adhesion (phospho-FAK) and the mammalian target-of-rapamycin (mTOR) pathway (phospho-mTOR and phospho-S6) in epidermal renewal and differentiation (Figure 3a and S10). Plotting the PC2-scores versus the measured phenotypes (Figure 3b-d), revealed that high-PC2 probes indeed have increased differentiation markers (Figure 3b), but also have increased cell-cycle arrest markers (Figure 3c). Indeed, keratinocyte differentiation is associated with a G2/M cell cycle arrest *in vivo* and *vitro*^29–31^. Strikingly, all high PC2 probes have a strong down-regulation of the mTOR pathway activity, suggesting an integral role in epidermal biology (Figure 3d). Thus, the ID-seq dataset captures relevant molecular phenotypes associated with keratinocyte differentiation.

**Figure 3.**
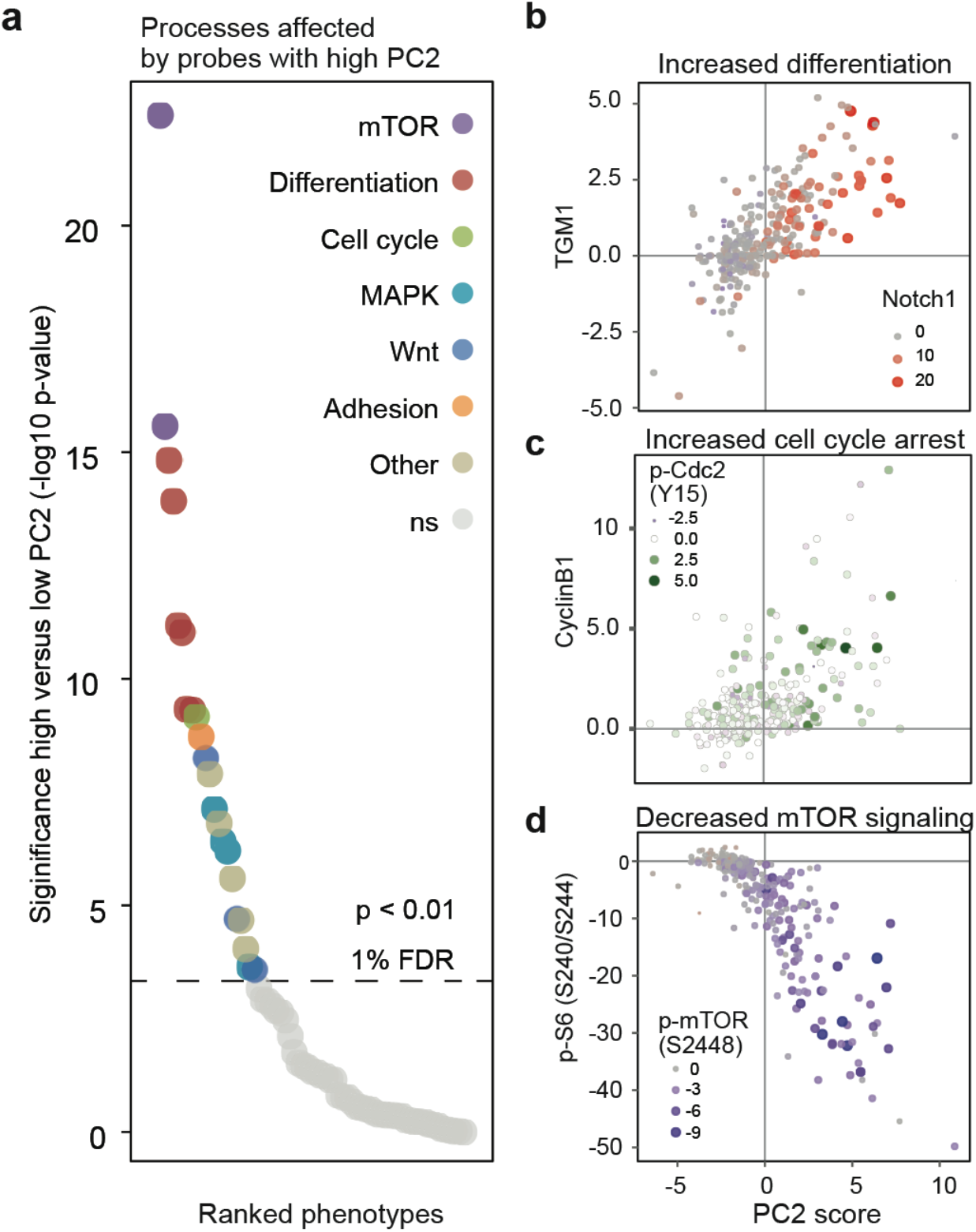
Probes with high PC2 affect phenotypes involved in differentiation, cell cycle arrest and mTOR signalling activity. (**a**) Summary of significantly (p < 0.01, FDR 1%, t-test, Figure S10) affected phenotypes from probes with high PC2 (top 10%) compared to low PC2 (bottom 10%) (**b**) Scatterplot illustrating probes with high PC2 score have increased TGM1 and Notch1 levels measured by ID-seq. **(c)** Scatterplot illustrating probes with high PC2 score have increased cell cycle arrest marker (CyclinB1 and p-cdc2) levels measured by ID-seq. (**d**) Scatterplot showing the probes with high PC2 score strongly have decreased phospho- S6 and phospho-mTOR levels illustrating decreased mTOR signalling activity measured by ID-seq.

### Identification of novel kinases involved in epidermal stem cell renewal

We reasoned that the inhibited kinases that strongly associated with PC2 are likely involved in epidermal stem cell renewal, as their inhibition leads to increased differentiation and cell-cycle arrest. To determine which kinases are inhibited by probes with high PC2 scores, we made use of available data on the biochemical selectivity and potency of the PKIS compounds towards 225 individual kinases.^26^ We applied outlier statistics to assign a set of inhibitory probes to each of these 225 kinases (p < 0.01, supplementary table 2). Subsequent Gene Set Enrichment Analysis (GSEA), identified 13 probe-sets enriched (p < 0.01, 1% FDR) in PC2 (i.e., probes inducing cell differentiation and/or cell cycle arrest) and of which the corresponding kinase is expressed in keratinocytes (Figures 4, S12 and S13). Our analysis returned the EGFR as the top hit, reflecting its key importance in epidermal stem cell renewal *in vitro* and *in vivo*. The PKIS probe library was designed as a platform for lead discovery and to provide chemical scaffolds for further medicinal chemistry^24–26^. Consequently, each probe may inhibit more than one kinase (Figure S14a). Importantly, the probes that inhibit the EGFR are distinct from those inhibiting the other kinases, indicating that the identification of these 12 kinases did not result from cross-reactivity of the probes towards the EGFR (Figure S14b). The list of other identified kinases included PRKD3 and FYN, two intracellular kinases shown to impact epidermal biology^32–36^. Moreover, immunohistochemical staining of human skin sections showed that the expression of the EGFR, RSK1, and PHKG1 is restricted to cells residing in the epidermal stem cell niche, whereas NUAK1 is expressed throughout the epidermis (Figure S15), consistent with our findings that inhibition of these kinases affects epidermal stem cell biology. Taken together, ID-seq identified both known and previously unrecognized kinase effectors of epidermal renewal across four major kinase families (Figure 2a, yellow nodes).

**Figure 4.**
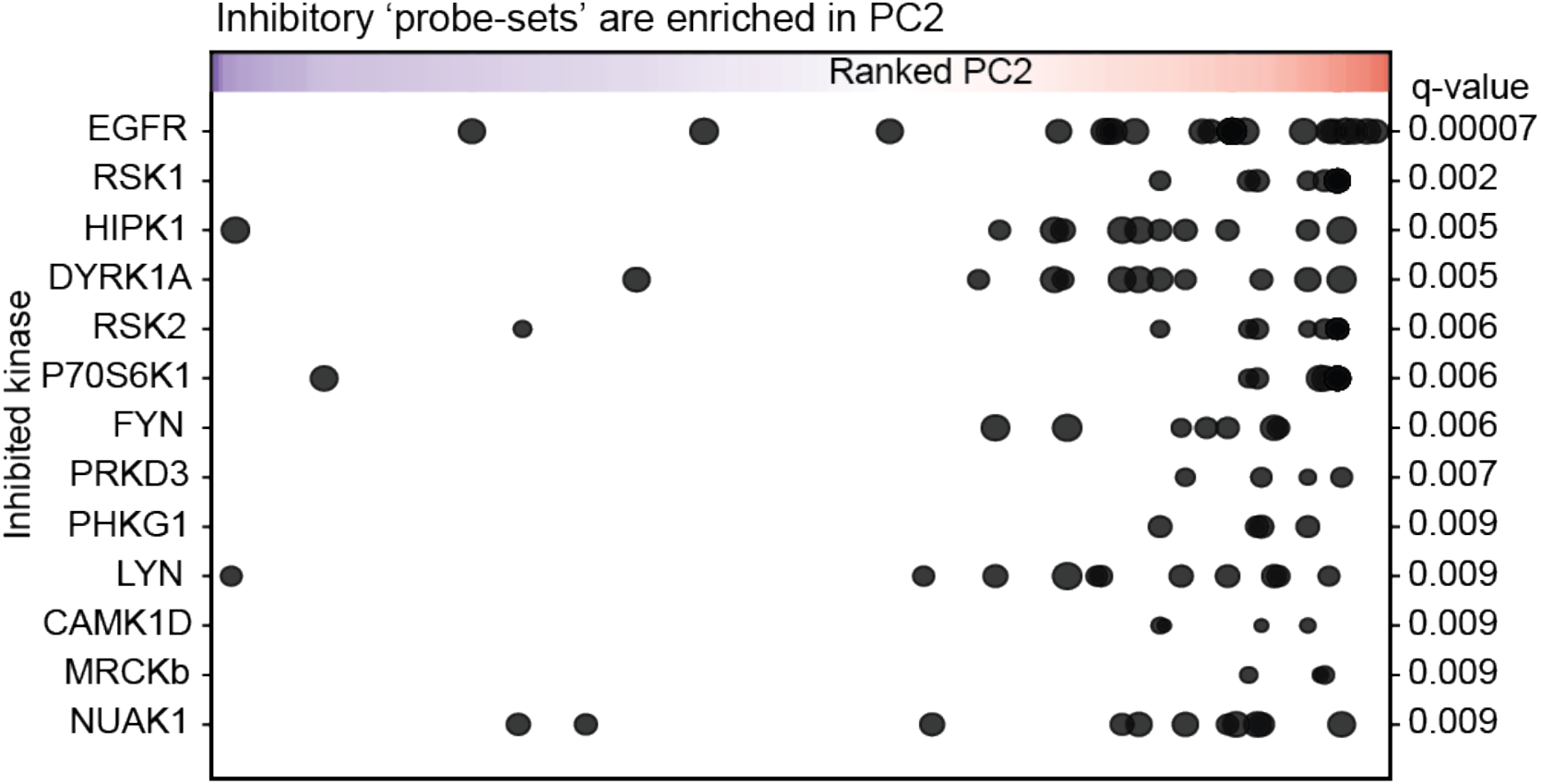
Gene Set Enrichment Analysis identifies kinases targeted by probes leading to epidermal differentiation. Summary plot of gene set enrichment analysis of probes in PC2 shows expressed inhibited kinases ordered according to the significance of enrichment (p-values (FDR) < 0.01, Figure S12). Probes (points in the graph) are ranked according to PC2 (x-axis), and point size shows % of inhibition for the indicated kinase by the probe.

### Four subclasses of kinases affect epidermal stem cells through distinct mechanisms

The rich information contained in the ID-seq dataset may be used to explore the underlying molecular mechanism of the kinase inhibition. For each kinase-set, we calculated the mean effect on each of the measured molecular phenotypes. These kinase-set level molecular profiles were used for both hierarchical clustering and PCA analysis, separating these 13 enriched kinases into 4 distinct subgroups (Figure 5a and b). The EGFR and its immediate downstream kinases, LYN and FYN, form a tight cluster, indicating that this grouping reflects molecular mechanistic relationships. In turn, this predicts that the kinases in the other subgroups may function through mechanisms that are different and potentially independent from the EGFR. If this were indeed the case, we would expect inhibition of these kinases to result in distinctive effects on (subsets of) the interrogated molecular phenotypes. We compared the inhibitory effects (average model derived estimate +/- SEM) on the 20 phenotypes distinguishing differentiated and renewing epidermal cells (as defined in Figure 2c) for exemplars of these 4 subgroups of kinases. Ranking these phenotypes based on the effect of EGFR inhibition and plotting the data of the other kinases in the same order showed that the overall trend in the measured phenotypes was conserved among the kinases, reflecting that these probe-sets indeed affect epidermal cell differentiation (figure 5c). Importantly, most of the kinases showed statistically significant deviations (p < 0.05) in distinct subsets of phenotypes compared to the EGFR, indicating that they function through discrete mechanisms to regulate epidermal renewal (Figure 5c). Together, this analysis shows that ID-seq allows highly multiplexed screening of hundreds of chemical probes, identifying kinases involved in epidermal renewal and at the same time provides information on the underlying molecular mechanism to categorize the identified effector kinases.

**Figure 5.**
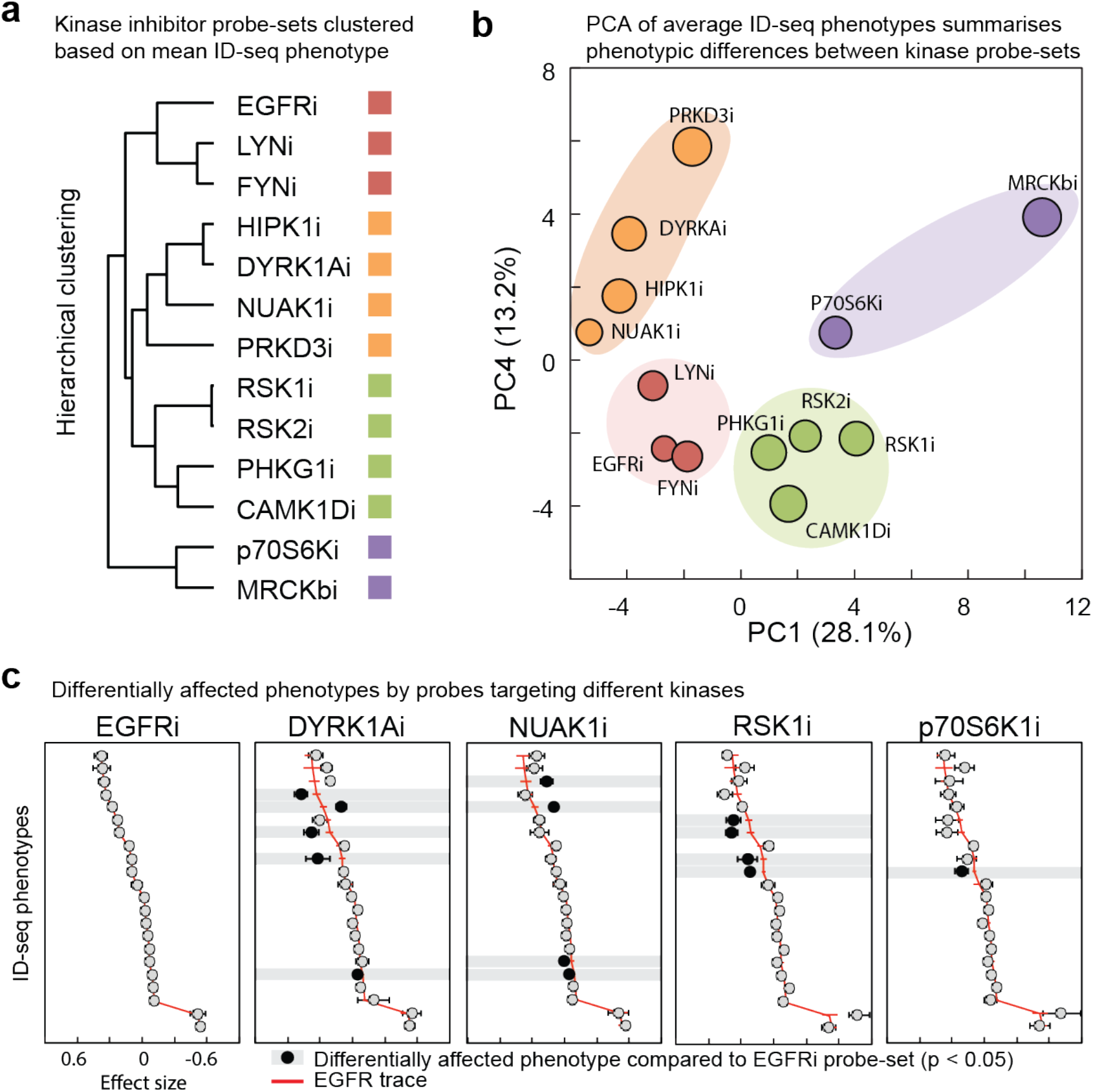
Enriched kinase inhibitor probe-sets show differentially affected molecular phenotypes. (**a**) K-means clustering of mean probe effect on phenotypes per kinase probe-set. (**b**) principal component analysis on mean probe effects shows in PC1 and PC4 distinct kinase probe-sets clusters. Colors based on K-means clustering of the data (see a) (**c**) The mean and standard error of probe effect on phenotypes, per probe-set inhibiting kinases EGFR, DYRK1A, NUAK1, RSK1/2/3/4, p70-Ribosomal S6 kinase. Compared to EGFR the other four kinases have a comparable phenotypic profile with several changes different effects on phenotypes (black nodes, p < 0.05, t-test).

### Selective inhibition of DYRK1A, NUAK1, RSK1-4 and p70S6K reduces epidermal stem cell colony forming potential

Finally, to verify the importance of the kinases representing these 4 subgroups in epidermal self-renewal, we obtained proven highly selective inhibitors of the EGFR, RSK1-4, p70S6K, NUAK1, and DYRK1A and examined their effect on epidermal stem cell function using *in vitro* colony formation assays (Figure 6a and b). For this, stem cells were first allowed to adhere to the culture plate containing a layer of feeder cells. Twenty-four hours after seeding the cultures were exposed to the kinase inhibitors in a broad range of concentrations for the remainder of the culture period. Subsequent automated imaging based quantification of the resulting colonies revealed that inhibition of the EGFR, RSK1-4, and DYRK1A decreased the self-renewal capacity of the epidermal stem cells (as determined by colony size) and stimulated the expression of the late differentiation marker TGM1. In contrast, p70S6K and NUAK1 inhibition resulted in decreased renewal but did not increase TGM1 expression. These results highlight the importance of these kinases in epidermal renewal.

**Figure 6.**
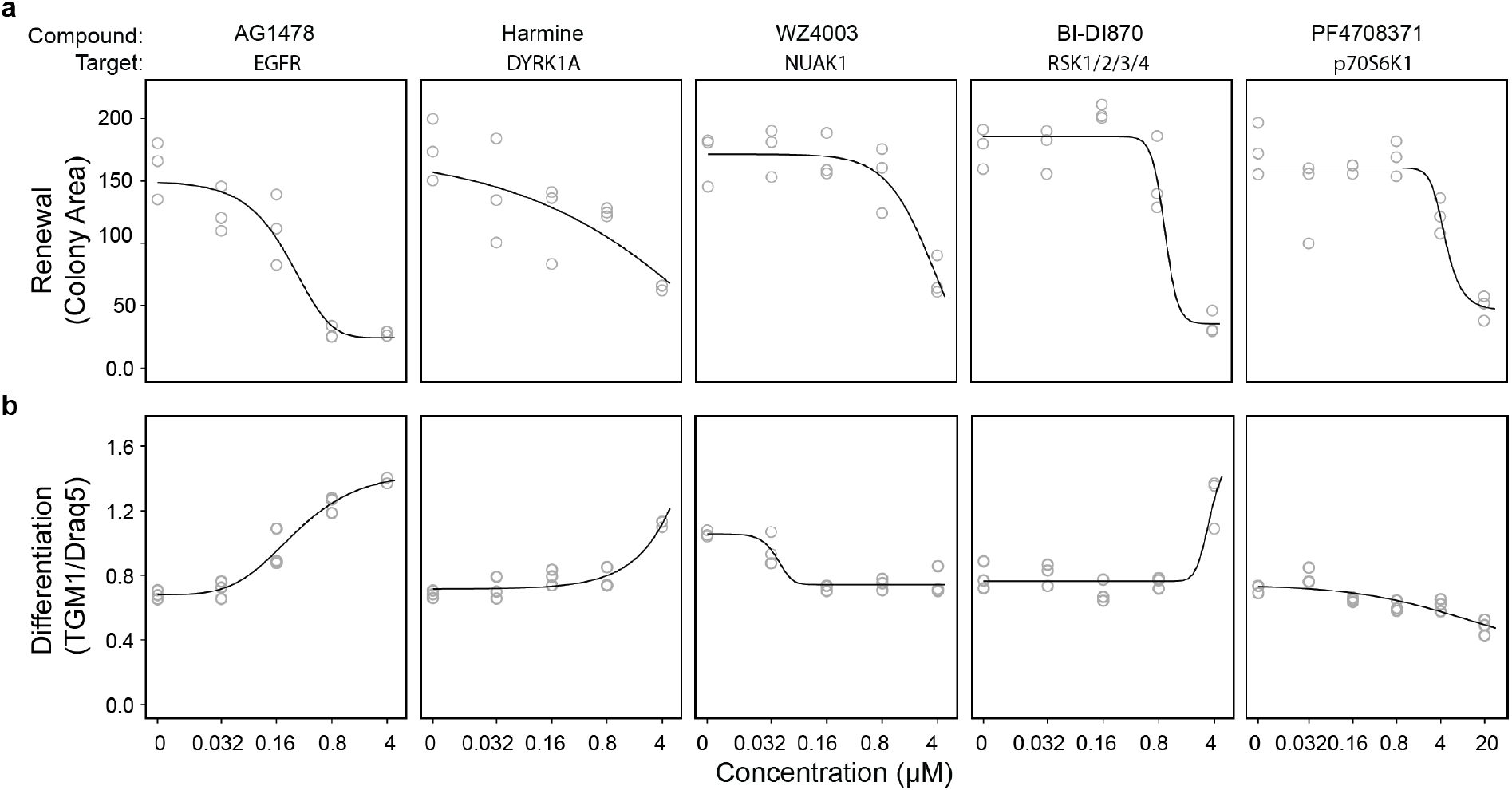
Renewal assay confirms crucial role for DYRK1A, NUAK1, RSK and p70S6 kinases in skin stem cell renewal while the inhibitors have distinct effect on late differentiation marker TGM1. (**a**) Colony area of keratinocytes (mean area of all colonies per replicate) after nine days of growth in the presence of specific kinase inhibitors AG1478, Harmine, WZ4003, BI-DI870 and PF4708371 targeting EGFR, DYRK1A, NUAK1, RSK1/2/3/4, and p70-Ribosomal S6 kinase respectively. Line shows modelled dose-response curves based on 3 biological replicates per concentration. (**b**) Levels of late differentiation marker (mean of all colonies per replicate), determined by immuno-fluorescent staining of TGM1 in CFA cell populations (corrected for cell number using Draq5 staining). Modelled dose-response curves (n=3) shows increase in TGM1 levels for 3 out of 5 inhibitors.

## Discussion

Cell-based phenotypic screens are frequently used in academia and the pharmaceutical industry to identify leads for drug development^37^,^38^. However, obtaining insight into the molecular mechanism of action of the selected compound is generally time consuming and expensive^39–41^. We developed the Immuno-Detection by sequencing (ID-seq) technology as an approach to facilitate high throughput highly multiplexed molecular phenotyping. We showed that ID-seq is precise, sensitive and applicable to large numbers of samples. We applied ID-seq on primary human epidermal keratinocytes in conjunction with the Published Kinase Inhibitor Set (PKIS) and identified 13 kinases that are important for epidermal stem cell function. Since many of the targeted therapies that are approved or in clinical trials are directed against kinases^41^,^42^, our findings serve as an important resource to identify potential drugs that could lead to severe skin toxicity. The straightforward ID-seq workflow was designed to be compatible with automation for applications in industry and will enable many potential mechanisms of action to be assayed for 100s, potentially even 1000s of compounds in a single experiment. In principle, ID-seq can be applied to any cell system, any perturbation and with any validated high-quality antibody, making it a flexible solution for large-scale high-dimensional phenotyping.

## Acknowledgements

We would like to thank E. Janssen-Megens and S. van Genesen for technical support and discussions, and Henk Stunnenberg, Ana Pombo and Michiel Vermeulen for discussions and critical reading of the manuscript. We would like to thank Bill Zuercher for discussions and reagents. The PKIS was supplied by GlaxoSmithKline, LLC and the Structural Genomics Consortium under an open access Material Transfer and Trust Agreement: http://www.sgc-unc.org. K.M., J.v.B, J.G., M.H and S.T are financially supported by by the Radboud University, the Dutch Organisation for Scientific Research (NWO-VIDI) and the European Union (Marie-Curie Career Integration Grant).

## Author contributions

J.v.B. designed the experiments, designed ID-seq barcodes, performed the ID-seq experiments and colony formation assays, wrote the code and R-package, analysed the data. J.G. and M.H. performed the iPCR and RNA-seq experiments, J.M. and R.v.d.S. produced all oligo DNA nucleotides, C.A developed the model for ID-seq count data analysis, K.M. conceived and oversaw the study and analysed the data, K.M. and J.v.B. wrote the manuscript with input from all other authors.

## Data availability

Used antibodies and oligo sequences are available as Supplementary Table 1 and Supplementary Table 3, respectively. Sequencing data and processed data from ID-seq experiments is available through GEO^43^ Series accession number GSE100135 https://www.ncbi.nlm.nih.gov/geo/query/acc.cgi?acc=GSE100135)

## Materials and Methods

### Cell culture

Keratinocytes (pooled foreskin strain KNP, Lonza) were expanded as described^31^ supplemented with Rock inhibitor (Y-27632, 10 µM). After expansion of the keratinocytes on feeders, the cells were grown for 1-3 days on keratinocyte serum-free medium (KSFM) with supplements (bovine pituitary extract (30 µg/mL) and EGF (0.2 ng/mL, Gibco) in a 96 or 384 wells plate at ~10.000 cells per well. Before 48 hours AG1478 treatment (10 µM), the cells were cultured for 48 hours in a 96 wells plate. Before EGF stimulation (100 ng/ml), keratinocytes were grown for three days on KSFM with and one day without supplements. After starvation, every 5 minutes EGF was added (n = 8). For the PKIS screen, 10,000 cells were seeded in a 384 wells plate and grown for 24 hours in KSFM with supplements, followed by 24-hour treatment with PKIS compounds (10 µM) or DMSO.

### Conjugation of antibodies to dsDNA

Antibodies and dsDNA were functionalised and conjugated as described^44^. See Supplementary Table 1 for a list of antibodies. In short, antibodies were functionalized with NHS-s-s-PEG4-tetrazine in a ratio of 1:10 in 50 mM borate buffered Saline pH 8.4 (150 mM NaCl). Then, N3-dsDNA was produced and functionalized with DBCO-PEG12-TCO (Jena Bioscience). See Supplementary Table 3 for a list of oligo sequences. Finally, purified functionalized antibodies were conjugated to purified functionalized DNA by 4-hour incubation at room temperature in borate buffered saline pH 8.4 in a ratio of 4:1 respectively. The reaction was quenched with an excess of 3,6-diphenyl tetrazine. After pooling, the conjugates were incubated with ProtA/G beads in BBS overnight. After thorough washes with PBS, the conjugates were eluted from the beads with 0.1 M citrate pH2.3 and immediately neutralized with TrisHCl pH 8.8. Subsequently, a buffer exchange into PBS (pH7.4) was performed using two Zeba-spin desalting columns. The size of the eluted DNA was checked on an agarose gel, confirming that all unconjugated DNA was removed using this purification approach.

### Antibody characterization

Detailed information on the antibodies is summarized in Supplementary Table 1. In brief, we selected antibodies suitable for immuno-histochemistry or immunofluorescence validated by manufacturer. The DSHB antibodies were produced and purified as described^44^. We performed antibody-dilution series with all antibodies on our primary human keratincoytes to show antibody-dependent signal via immuno-fluorescence. Moreover, we coupled antibodies to DNA barcodes as described in our conjugation and immuno-PCR protocol^44^. These antibodies show antibody concentration-dependent signal via immuno-PCR, indicating successful conjugation, release and detection of DNA-tags. Finally, we performed several modulation experiments to show specific dynamics in our keratinocytes measured via the antibodies in immunofluorescence, immuno-PCR (Supplementary Table 1) or ID-seq (Figure S5).

### Immuno-staining with Antibody-DNA conjugates and release of DNA tags

Keratinocytes were fixed with 4% paraformaldehyde in PBS for 15 minutes at RT, washed three times with PBS and stored at 4 °C (up to 3 - 4 days before further use). Then, cells were permeabilized and blocked for 30 minutes using 0.5x protein free blocking buffer (Thermo Fisher) in PBS with 0.1 % Triton and 200 ng/ml single strand Salmon Sperm DNA (sssDNA). Blocking the cells and wells with sssDNA is crucial to suppress background binding of the Ab-DNA conjugates^44^. Then, cells were incubated with conjugated antibodies in the same buffer at 0.1 µg/mL antibody, at 4 °C overnight. After immunostaining with conjugates, the cells were thoroughly washed with PBS (3x short, 3x 15 min, 3x short). Then, release buffer was freshly prepared (10 mM DTT in borate buffered saline pH 8.4). Cells were incubated with 20 to 50 µl of release buffer depending on the plate type and well-size and incubated for 90 minutes at RT, with careful mixing (on a vortex) every 30 minutes. Released DNA barcodes were collected and stored at −20 °C.

### Sample barcoding and sequencing library preparation

To barcode the released DNA-tags from each cell population (See Supplementary Note 1 for sequence design), a 25 µl PCR was performed per sample containing 8-15 µl sample with released DNA-tags, 0.2 mM dNTPs, 1 ul PFU polymerase, 1x PFU buffer (20mM Tris-HCl pH8.8, 2mM MgSO_4_, 10mM KCl, 10 mM (NH_4_)_2_SO_4_, 0.1% triton, 0.1 mg/ml BSA), spike-in DNA barcodes, forward primer (AATGATACGGCGACCACCG, Biolegio) and a well specific reverse primer (Supplementary Table 3, Supplementary Scheme 1b). In a 96 wells PCR machine (T100 Thermal Cycler, Biorad) the following program was used: 1) 3 min at 95 °C, 2) 30 sec at 95 °C, 3) 30 sec at 60 or 54 °C, 4) 30 sec at 72 °C, 5) repeat 2-4 nine times, 6) 5 min at 72 °C, 7) ∞ 12 °C. Then, all well-specific labelled DNA barcodes from one plate were pooled to 1 sample. This sample was purified using a PCR purification column (Qiagen) according to manufacturer’s protocol. Samples were eluted with 30 µl nuclease-free water. To remove any residual primers, samples were treated with Exonuclease I in 1x PFU buffer for 30 min at 37 °C. After inactivation for 20 min at 80 °C, another 25 µl PCR reaction was prepared with 15 - 17 µl sample, 0.2 mM dNTPs, x 1 µl PFU polymerase, 1x PFU buffer, forward primer (AATGATACGGCGACCACCG, Biolegio) and a sample specific reverse primer with Illumina index barcode and adapter sequence (Supplementary Table 3, Supplementary Scheme 1d). The same program as PCR reaction (I) was used, and reactions were purified over a PCR purification column (Qiagen). All PCR reactions were then incubated for 45 minutes with one µl Exonuclease I to remove residual primers. The PKIS screen samples were further size-selected using size selection columns (Zymo, according to manufacturer’s protocol) for fragments > 150 bp. Finally, all samples were purified over PCR purification mini-elute column (Qiagen) and eluted in 10 µl elution buffer. Final sequencing samples were run on a 2% agarose gel (0.5x TBE) with 10× SYBR Green I (Life Technologies) and scanned on a Typhoon Trio+ machine (GE Healthcare), or analysed with the 2100 Bioanalyzer (Agilent) to confirm the size of the DNA fragments (expected size around 185 bp).

### ID-seq data analysis

Sequence data from the NextSeq500 (Illumina) was demultiplexed using bcl2fastq software (Illumina). The quality of the sequencing data was evaluated using a FastQC tool (version 0.11.4 and 0.11.5, Babraham Bioinformatics). Then, all reads were processed using our dedicated R-package (IDSeq, Supplementary Note 2). In short, the sequencing reads were split using a common “anchor sequence” identifying the position of the UMI sequence, Barcode 1 (antibody specific) and Barcode 2 (well specific) sequence. After removing all duplicate reads, the number of UMI sequences were counted per barcode 1 and 2. Finally, barcode 1 and barcode 2 sequences were matched to the corresponding antibody and well information.

Using R-package *DESeq2*^45^, we calculated normalisation factors (Estimated Size factor) to account for differences in sequencing depth per sample. Using *lme4*, we analysed the effect of a specific condition using a linear mixed effect model (Supplementary Note 3).

For each antibody in the PKIS screen the effect and significance of each treatment was determined as described in Supplementary Note 3. Then the ‘signed p-value’ was derived from the sign of the model estimate (positive/negative) and the p-value. This signed p-value was used as input for the PCA analyses (Figure 2b and S9). To calculate effects of probe-sets per phenotype, the mean model estimate was calculated. These means were used for subsequent PCA analysis and ‘phenotype profiles’ described in Figure 5.

### Immuno-PCR experiments

The immuno-PCR experiments were performed as described previously in our paper on antibody-DNA conjugates^44^. In short, each antibody was conjugated to dsDNA, and used in an immunostaining as described. DNA was released using 10 mM DTT in BBS pH8.4 and measured by quantitative PCR using iQTM SYBR Green Supermix on CFX 96 machine. The 2^-Ct values were used to calculate the mean signal and standard deviation from 4 biological replicates. The Pearson correlation between these immuno-PCR and the multiplexed ID-seq signal was calculated using the mean.

### RNA expression levels of kinases using CEL-seq2 mRNA quantification

mRNA sequencing was performed according to the CELseq2 protocol^46^ with adaptations. Reverse transcription was performed in 2 µl reactions overlaid with 7 µl Vapor-Lock (Qiagen) using Maxima H minus reverse transcriptase (ThermoFisher) and 100 pg purified RNA per sample. Primer sequences were adapted to allow sequencing of 63 nucleotides of mRNA in read 1 and 14 nucleotides in read 2, comprising the sample barcode and UMI. Reverse transcription primer: 5’GCCGGTAATACGACTCACT-ATAGGGGTTCAGACGTGTGCTCTTCCGATCTNNNNNNNN[6ntsamplebarcode]TTTTTTTTTTTTTTTTTTTTT TTTV3’, random-octamer-primer for reverse transcription of amplified RNA: 5’CACGACGCTCTTCCGATCT-NNNNNNNN3’, library PCR Primers: 5’AATGATACGGCGACCACCGAGATCTACACTCTTTCCCTACACG-ACGCTCTTCCGATCT3’ and 5’CAAGCAGAAGACGGCATACGAGAT[6ntindex]GTGACTGGAGTTCAGACGTGTG-CTCTTCCGATC3’. Sequencing was performed using the NextSeq500 from Illumina.

### Colony formation assay

In 6 wells plate, 200.000 feeders (J2-3T3) cells were seeded in DMEM (with 10% BS and 1% pen/strep). After one day, feeder cells were inactivated by 3-hour treatment with mitomycin C. After thorough washes with DMEM, 1000 keratinocytes were seeded into each well^31^. The following day, treatment was started (day 0) by refreshing medium and addition of the indicated concentration of compound, or DMSO as a vehicle control. Cells were grown in the presence of compounds for eight more days, and the medium was refreshed on days 2 and 5. Rocki was present until day 2 of the treatment. Cells were fixed, stained with TGM1 specific antibodies and scanned as described before ^27^. Raw images from the LiCor Odessey system were processed with CC Photoshop and CellProfiler with consistent settings. Data obtained via automatic counting and imaging analysis via CellProfiler was analysed and visualized in the R programming language.

